# Neuronal kinase SGK1.1 protects against brain damage after status epilepticus

**DOI:** 10.1101/2020.04.03.024067

**Authors:** Elva Martin-Batista, Laura E. Maglio, Natalia Armas-Capote, Guadalberto Hernandez, Diego Alvarez de la Rosa, Teresa Giraldez

## Abstract

Epilepsy is a neurological condition associated to significant brain damage produced by *status epilepticus* (SE) including neurodegeneration, gliosis and ectopic neurogenesis. Reduction of these processes constitutes a useful strategy to improve recovery and ameliorate negative outcomes after an initial insult. SGK1.1, the neuronal isoform of the serum and glucocorticoids-regulated kinase 1 (SGK1), has been shown to increase M-current density in neurons, leading to reduced excitability and protection against seizures. We now show that SGK1.1 activation potently reduces levels of neuronal death and gliosis after SE induced by kainate, even in the context of high seizure activity. This neuroprotective effect is not exclusively a secondary effect of M-current activation but is also directly linked to decreased apoptosis levels through regulation of Bim and Bcl-x_L_ cellular levels. Our results demonstrate that this newly described antiapoptotic role of SGK1.1 activation acts synergistically with the regulation of cellular excitability, resulting in a significant reduction of SE-induced brain damage. The protective role of SGK1.1 occurs without altering basal neurogenesis in brain areas relevant to epileptogenesis.

**SIGNIFICANCE STATEMENT:** Approaches to control neuronal death and inflammation are of increasing interest in managing epilepsy, one of the most important idiopathic brain diseases. We have previously shown that activation of SGK1.1 reduces neuronal excitability by increasing M-current levels, significantly reducing seizure severity. We now describe a potent neuroprotective role of SGK1.1, which dramatically reduces neuronal death and gliosis after status epilepticus. This effect is partially dependent on M-current activation and includes an additional anti-apoptotic role of SGK1.1. Our data strongly support the relevance of this kinase as a potential target for epilepsy treatment.

## INTRODUCTION

Epilepsy is a neurological disorder characterized by recurrent and unpredictable seizures, affecting more than 50 million people around the world. Temporal Lobe Epilepsy (TLE) is one of the most common types of epilepsy and the most refractory to treatment in humans. Two-thirds of TLE patients develop a sclerosis state caused by neuronal loss and gliosis (Mathern et al., 1997). Epileptogenesis has been associated to neurodegeneration (Dingledine et al., 2014), gliosis (Sofroniew, 2014), uncontrolled inflammation (Vezzani et al., 2011), and aberrant neurogenesis (Jessberger and Parent, 2015). This alteration in neurological homeostasis often results in a chronic epileptic condition (Pitkanen and Lukasiuk, 2009). Therefore, there is an increasing interest in the field of epilepsy research to develop new therapies targeting neuronal death and inflammation pathways.

*Status epilepticus* (SE) is defined as a period of continuous behavioral or electroencephalographic seizures lasting more than 30 min. Chemically-induced-SE animal models reproduce most behavioral, electroencephalographic and neuropathological characteristics of TLE and have been widely used in basic epilepsy research to study this disorder (Levesque et al., 2016). SE-triggered negative outcomes in the brain include neuronal death, which has been associated to the appearance of new seizures (Mathern et al., 2002). Over-activation of glutamate receptors following SE constitutes a major cause of neuronal death. Elevation of Na^+^ and Ca^2+^ influx eventually leads to membrane disruption, cell lysis, release of free radicals and DNA degradation (Wang et al., 2005; Fujikawa, 2006). In addition, alterations in the expression of various biochemical markers for apoptosis have been related to seizure-induced neuronal loss, including members of the Bcl-2 gene family and caspases (Kim et al., 2014). Therefore, neuroprotection could be facilitated by two different but not necessarily unrelated general mechanisms. First, the activation of pathways limiting excitability. Secondly, the prevention of downstream events leading to apoptosis.

We have recently shown that increased activity of SGK1.1, the neuronal isoform of the serum and glucocorticoids-regulated kinase 1 (SGK1) potently reduces seizure duration and severity in a kainic acid (KA)-induced SE paradigm with transgenic mice expressing a permanently active form of the kinase (Armas-Capote et al., 2019). Electrophysiological measurements in sympathetic neurons (Miranda et al., 2013) and hippocampal brain slices (Armas-Capote et al., 2019) showed that SGK1.1 up-regulates the neuronal M-current mediated by Kv7.2/7.3 ion channels. Altogether, our data suggested that SGK1.1 activation provides a mechanism securing the brain against epilepsy-related neuronal damage by controlling neuronal excitability.

In addition to the control of neuronal excitability, different studies have demonstrated that seizure-induced neuronal loss can be diminished by molecular and pharmacological approaches modulating apoptotic cell death (Henshall et al., 2002; Roy et al., 2002). Interestingly, SGK1 isoforms share high homology to the catalytic domain of the well described anti-apoptotic kinase AKT (Kobayashi et al., 1999). Both AKT and its upstream activator PI3K have been demonstrated to reduce apoptosis and promote neuronal survival in the central nervous system (Datta et al., 1997; Crowder and Freeman, 1998). The ubiquitous isoform SGK1 shows cell-survival and anti-apoptotic effects (Brunet et al., 2001; Ferrelli et al., 2015) and has been linked to neuroprotection in stroke (McCaig, 2019; Wang, 2019). Taking all this into account, we hypothesized that in addition to the observed reduction in neuronal excitability, SGK1.1 activation could directly reduce apoptosis levels post-SE. In this study we have extensively quantified neuronal death levels following KA-induced SE in different brain areas of WT and transgenic mice with constitutively active SGK1.1 expression. Our data show that increased SGK1.1 activity in the transgenic mouse model dramatically reduces neuronal death associated to KA-induced seizures. This effect is associated to significantly diminished inflammatory processes and the absence of ectopic neurogenesis. Finally, activation of the kinase is associated to altered levels of apoptotic biomarkers consistent with lower apoptosis levels. In summary, we demonstrate the existence of a dual mechanism underlying SGK1.1-mediated protection against neuronal death involving not only the modulation of neuronal excitability, but also the additional modulation of cellular pathways leading to apoptosis.

## MATERIAL AND METHODS

### Antibodies and chemicals

Antibodies against Bcl-2 interacting mediator of cell death (Bim;sc-374358) and B-cell lymphoma-extra large (Bcl-x_L_;sc-8392) were obtained from Santa Cruz Biotechnology (Santa Cruz, CA); anti-parvalbumin (PV;P3088) and anti-glial fibrillary acidic protein (GFAP;G3893) antibodies were obtained from Sigma-Aldrich; bromodeoxyuridine (BrdU, NBP2-14890) was obtained from Novus; rabbit Anti-Goat IgG conjugated to Alexa Fluor® 488 (ab150141), anti-Doublecortin (DCX;ab153668) and anti-ionized calcium-binding adapter molecule 1 (Iba-1;ab5076) were obtained from Abcam; goat anti-mouse IgG conjugated to Alexa Fluor® 594 (A-11005), goat anti-chicken IgY conjutated to Alexa Fluor® 594 (A-11042), goat anti-rabbit IgG conjugated to Alexa Fluor® 488 (A-11008) and anti-calcium/calmodulin-dependent protein kinase II (CaMKII;MA1-048) were obtained from Thermo Fisher Scientific. A rabbit polyclonal antibody against SGK1.1 was generated against a glutathione S-transferase (GST) fusion protein containing amino acids 41-80 from the SGK1.1-specific exon 2, expressed in *E. coli* and purified by affinity chromatography. KA (K0250) and M-current blocker XE991 (X2254) were obtained from Sigma Aldrich. SGK1 inhibitor EMD638683 was from MedChem Express (HY-15193A).

### Animals and seizure induction

Animal handling and experimental procedures were approved by Universidad de La Laguna Ethics Committee and conform to Spanish and European guidelines for protection of experimental animals (RD53/2013; 2010/63/EU). This work is based on the use of a transgenic mouse model previously generated in our laboratory, B6.Tg.sgk1 (Miranda et al., 2013; Armas-Capote et al., 2019). Briefly, the transgenic line was created with a bacterial artificial chromosome containing the whole Sgk1 gene and a point mutation (S422D in the ubiquitous isoform SGK1; S515D in the neuronal isoform SGK1.1) that renders the kinase constitutively active. The BAC expresses Sgk1 splice isoforms under their own promoters and results in expression of the neuronal isoform SGK1.1 in the brain (Arteaga et al. 2008). Founder animals were crossed with wild type C57BL/6J for at least nine generations. All experiments described here were performed in homozygous mice obtained by crossing heterozygous animals. FVB.Tg.sgk1 mice were generated by transferring the transgene to the inbred strain FVB/N, backcrossing for 9 generations (Armas-Capote et al., 2019). Wild type mice (B6.WT or FVB.WT) in the same genetic background (C57BL/6J or FVB/N) were used as controls. Seizures were induced in 16-22 weeks-old mice with intraperitoneal (ip) injection of 20 mg/kg KA as previously described (Armas-Capote et al., 2019). Seizure behavior was evaluated for 2 h using a modified Racine scale: 0, no response, 1, immobility, 2, rigid posture and tail extension, 3, head bobbing and repetitive movements, 4, rearing, 5 rearing and falling continuously, 6, tonic-clonic generalized seizures and 7, death. Only mice reaching score 6 were included in the histological studies. Where indicated, XE991 (10 mg/kg) or EMD638683 (1.6 mg/kg) were administered ip 1 h or 1 day prior to KA injection respectively, as described elsewhere (Armas-Capote et al., 2019). Mice injected with saline were used as controls.

### Western Blot Analysis

Hippocampi were collected 24 h after KA treatment and protein extracts were obtained by processing samples with lysis buffer (Tris-Base 0.5 M, pH 7.4; SDS 10 %) containing phosphatase and proteinase inhibitors (Roche). Lysates were clarified by centrifugation at 14000 x g for 10 min and protein concentration was measured using the bicinchoninic acid assay. Proteins were separated by SDS denaturing gel electrophoresis (SDS-PAGE), transferred to a polyvinylidene fluoride membrane and analyzed by western blot (WB). Chemiluminescence signals were analyzed using Image Lab® software 6.0 (Bio-Rad).

### Histological sample preparation

Three days after KA treatment, mice were deeply anesthetized (sodium pentobarbital, 40 mg/kg, ip) and perfused transcardially with saline (NaCl 0,9%) and paraformaldehyde (PFA) 4% in 0.1 M phosphate-buffered saline pH 7.4 (PBS). Dissected brains were post-fixed in PFA for 24 h (4°C) before being transferred to sucrose 30% for cryoprotection overnight (−80°C). Thick coronal sections (30 μm) were obtained using a freezing microtome. Saline-injected animals were used as controls.

### Neurodegeneration quantification by Fluoro-Jade C staining

Coronal slices were stained with Fluoro-Jade C (FJC), an anionic fluorochrome capable of selectively staining degenerating neurons in brain slices (Schmued et al., 1997). Neuronal degeneration was evaluated 72 h after KA injection in WT and Tg.sgk1 mice reaching Racine stage 6, following a previously described protocol (Afonso-Oramas et al., 2010). Sections were examined using confocal microscopy (Leica TCS SP8). Image analysis was performed using ImageJ software (Schindelin et al., 2012). Five different sections were selected for each animal (from −1.355 to −3.08 in reference to bregma, Allen Institute for Brain Science, 2014) providing a total of five individual values per region for each animal. Measurements from all sections were averaged to obtain one final measurement per region.

### Gliosis quantification by immunohistochemistry

The presence of reactive gliosis 72 h post-KA injection was assessed by measuring the expression of GFAP in astrocytes and Iba-1 in microglia. One set of coronal sections (between −1.755 and – 2.155 mm from bregma) was selected for evaluation of potential GFAP and Iba-1 variations after KA. Briefly, sections were washed three times with PBS, blocked with 4% goat serum for 1 h and incubated overnight with anti-GFAP at 1:400 or anti-Iba-1 at 1:500 diluted in 2% goat serum. Sections were then washed 3-4 times in PBS before incubation with secondary antibodies conjugated to Alexa Fluor® 594 at 1:200 or Alexa Fluor® 488 at 1:1000 for 2 h. Finally, sections were washed and mounted with Mowiol and 4′,6-diamidino-2-phenylindole (DAPI) for examination under confocal microscopy. Measurements from the two hemispheres were obtained and averaged as GFAP area/DAPI area using ImageJ software.

### Neurogenesis quantification by BrdU incorporation and immunohistochemistry

We tested the effect of SGK1.1 activation on ectopic neurogenesis processes using BrdU incorporation and DCX staining on brain slices. Adult B6.WT and B6.Tg.sgk1 BrdU-injected mice (50 mg/kg, twice) were perfused with PFA 4% 24 h after injection. Brains were removed and post-fixed in PFA 4% and 50 μm coronal brain slices were obtained with a freezing microtome. After blocking with 1% serum in PBS and 1% Triton X-100, and DNA denaturation with hydrochloric and boric acid (2 N at 37°C 30 min and 0,1M at RT 10 min respectively), proliferating cells were detected with rabbit anti-BrdU antibody at 1:500 (72 h at 4°C) followed by anti-rabbit Alexa Fluor® 488 antibody at 1:1000. Proliferating cells (BrdU^+^) in the hilus (ectopic neurogenesis) and subgranular layer (newborn cells) of the dentate gyrus (DG) were quantified and compared in transgenic and WT mice. Additionally, the amount of adult neuronal proliferation was evaluated by the presence of DCX-positive (DCX^+^) neuroblasts in DG using a chicken anti-DCX antibody at 1:500 and anti-chicken Alexa Fluor® 594 antibody at 1:1000 (Kuhn, 1996; Pérez-Domper et al., 2017).

### Cell culture and transfection

Human embryonic kidney cells HEK293T were obtained from the American Type Culture Collection (Manassas, VA) and maintained in DMEM supplemented with 10% FBS. Cells were transfected 24-48 h before experiments using Jetprime (Polyplus Transfection, Illkirch) following the manufacturer’s instructions. Plasmid constructs for WT, mutant SGK1.1 and AKT expression have been previously described (Coric et al., 2004; Arteaga et al., 2008; Wesch et al., 2010); pECFP-N1 was obtained from Clontech.

### Terminal deoxynucleotidyl transferase dUTP nick end labeling (TUNEL) Assay

In order to evaluate the potential anti-apoptotic role of SGK1.1, we performed TUNEL assays in HEK293T cells. Briefly, cells were transiently transfected with SGK1.1 dominant negative mutant (SGK1.1^K220A^), a mutant rendering the kinase constitutively active (SGK1.1^S515D^) or a mutant targeting it to the nucleus (SGK1.1^FF19,20AA^). AKT was used as a positive control for anti-apoptotic effects. An empty vector (pECFP-N1) was used to control for transfection toxicity. Cells were treated with 1 mM hydrogen peroxide (H_2_O_2_) for 4h to induce apoptosis. The TUNEL protocol was performed using a commercial kit (ApopTag® fluorescein in situ apoptosis detection kit, Millipore). Apoptotic cells were detected as localized bright green cells in a blue background (DAPI) by confocal microscopy.

### Experimental design and statistical analysis

Statistical analysis was performed using Prism 8 (GraphPad). Prior to any statistical analysis, we assessed normality distribution of our data using D’Agostino-Pearson or Shapiro-Wilk normality test. Once normality was established, we compared data means using t-test or one-way ANOVA test for parametric distributions and Mann-Whitney for non-parametric distributions. Specific statistical tests and number of experiments/samples and/or animals (indicated by n and N, respectively) are indicated in each figure legend.

## RESULTS

### SGK1.1 protects mice from kainate-induced neuronal death

We have previously shown that transgenic mice expressing a constitutively active form of the kinase (B6.Tg.sgk1) show significantly reduced seizure severity in the KA-induced SE animal paradigm (Armas-Capote et al., 2019). This is reflected in fewer mice reaching more severe Racine stages and associated mortality (Figure 1A). In this study we evaluated the impact of increased SGK1.1 activity in neurodegeneration and gliosis on mice surviving KA treatment. For this purpose, we selected mice reaching Racine stage 6 (generalized tonic-clonic seizures; Figure 1A, orange box). We reasoned that the comparison of transgenic mice with their WT counterparts that are reaching comparable seizure severity levels would allow us to unveil the potential contribution of additional neuroprotective mechanisms activated by SGK1.1 independent of seizure protection (Armas-Capote et al., 2019). Neurodegeneration was quantified by FJC staining of brain slices from B6.WT and B6.Tg.sgk1 mice reaching tonic-clonic seizures (Racine 6). As shown in Figure 1B (left panels), significant levels of neuronal damage were observed in B6.WT mice, reflected as a significant number of FJC-stained cells (Figure 1C). This result was comparable to previous reports using similar experimental conditions (Hopkins et al., 2000; Yan et al., 2018). In contrast, activation of SGK1.1 in B6.Tg.sgk1 neurons conferred a remarkable level of protection against neuronal death in all brain areas studied, including hippocampus, somatosensorial auditive and piriform-entorhinal cortex (Figure 1B, right panels, and Figure 1C). This observation is especially striking taking into account that all B6.Tg.sgk1 mice reached tonic-clonic convulsions, suggesting the possibility that protective mechanisms are activated by SGK1.1 even in conditions of neuronal hyperexcitability. The neuroprotective effect associated to SGK1.1 activation was not specific of the C57BL/6J genetic background. Similar results were obtained with transgenic mice generated on the FVB background, which has been proposed to display increased seizure susceptibility to KA (Schauwecker and Steward, 1997; McKhann et al., 2003; Kasugai et al., 2007) (Figure 1D-F).

**Figure 1.**
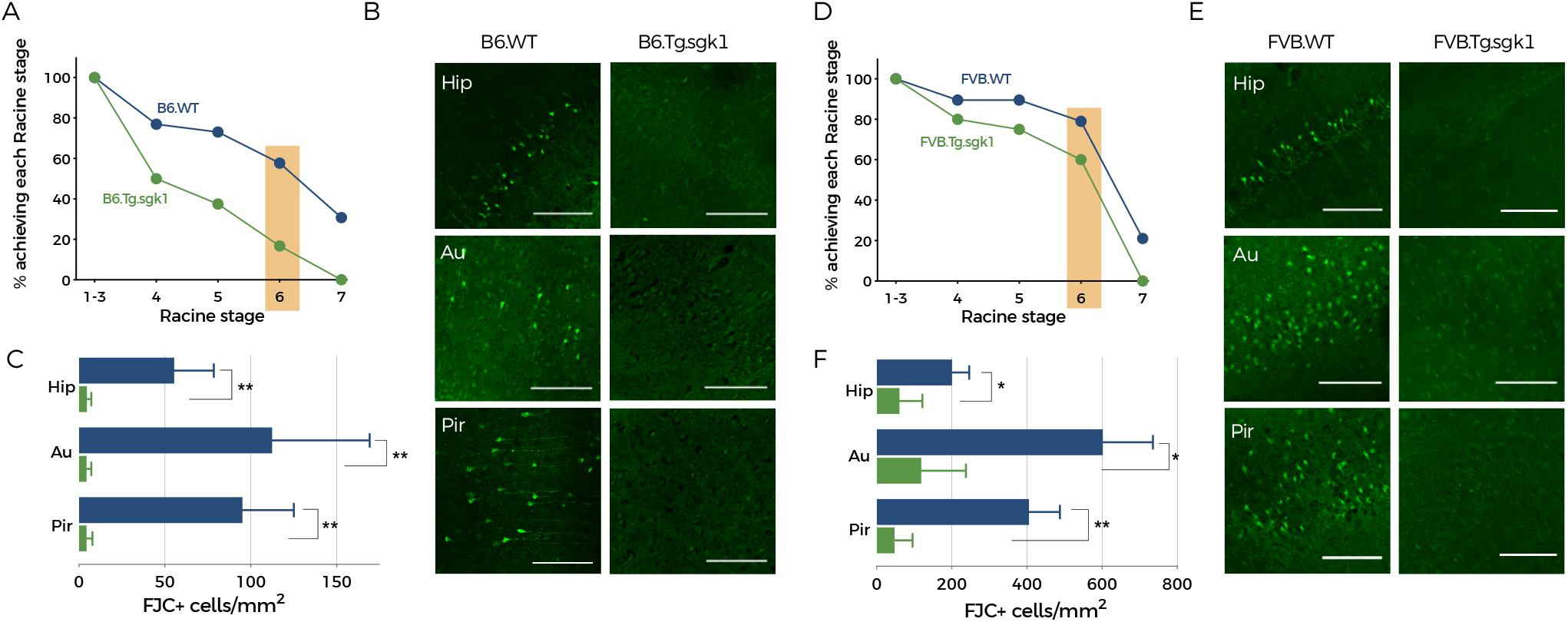
Fluoro-Jade C staining reveals significantly decreased neuronal in Tg.sgk1 mice after SE. (A) Cumulative plot representing the percentage of B6.WT (blue) and B6.Tg.sgk1 (green) mice reaching the indicated Racine stages after KA injection. Stage 7 corresponds to death associated to SE. Only mice reaching stage 6 were used to evaluate neuronal death (highlighted in orange) (B6.WT N=26, B6.Tg.sgk1 N=24). Part of the data in this graph have been obtained from (Armas-Capote et al., 2019). (B) Representative confocal images of FJC stained brain sections B6.WT (left column) and B6.Tg.sgk1 mice (right column) reaching Racine stage 6. Hippocampus (Hip), auditive cortex (Au) and piriform cortex (Pir) brain areas are shown. Scale bar = 100 μm. (C) Quantitative analysis of neurodegeneration events (FJC-positive cells) normalized to area (mm^2^) corresponding to regions shown in B from B6.WT and B6.Tg.sgk1 mice brain slices. Data are mean ± SEM (N=5; n=25; Mann-Whitney test, **p<0.01). (D) Cumulative plot representing the percentage of FVB mice reaching the indicated Racine stages. Only mice reaching stage 6 were used to evaluate neuronal death (highlighted in orange; 72h after KA treatment) (FVB.WT N=19; FVB.Tg.sgk1 N=19). (E) Representative confocal images corresponding to FJC staining of brain slices from same regions as B, from FVB.WT (left column) and FVB.Tg.sgk1 mice (right column) reaching Racine stage 6. Scale bar = 100 μm. (F) Quantification of FJC positive cells normalized to area (mm^2^) in both genotypes on the FVB genetic background. Data are mean ± SEM (N=7; n=35; Mann-Whitney test, *p<0.05, **p<0.01)

To assess that the observed effects were a direct consequence of kinase activation, mice were acutely pre-treated with the SGK1.1 inhibitor EMD638683. As shown in Figure 2A and consistent with previous reports (Armas-Capote et al., 2019), inhibition of the kinase reverted the protective effect and abolished all differences in seizure behavior between genotypes (Figure 2A). Importantly, identical levels of neuronal death were observed in B6.WT and B6.Tg.sgk1 mice (Figure 2B-C) in all brain areas. These results demonstrate that kinase activity underlies the mechanism reducing neuronal death associated to KA-induced seizures.

**Figure 2.**
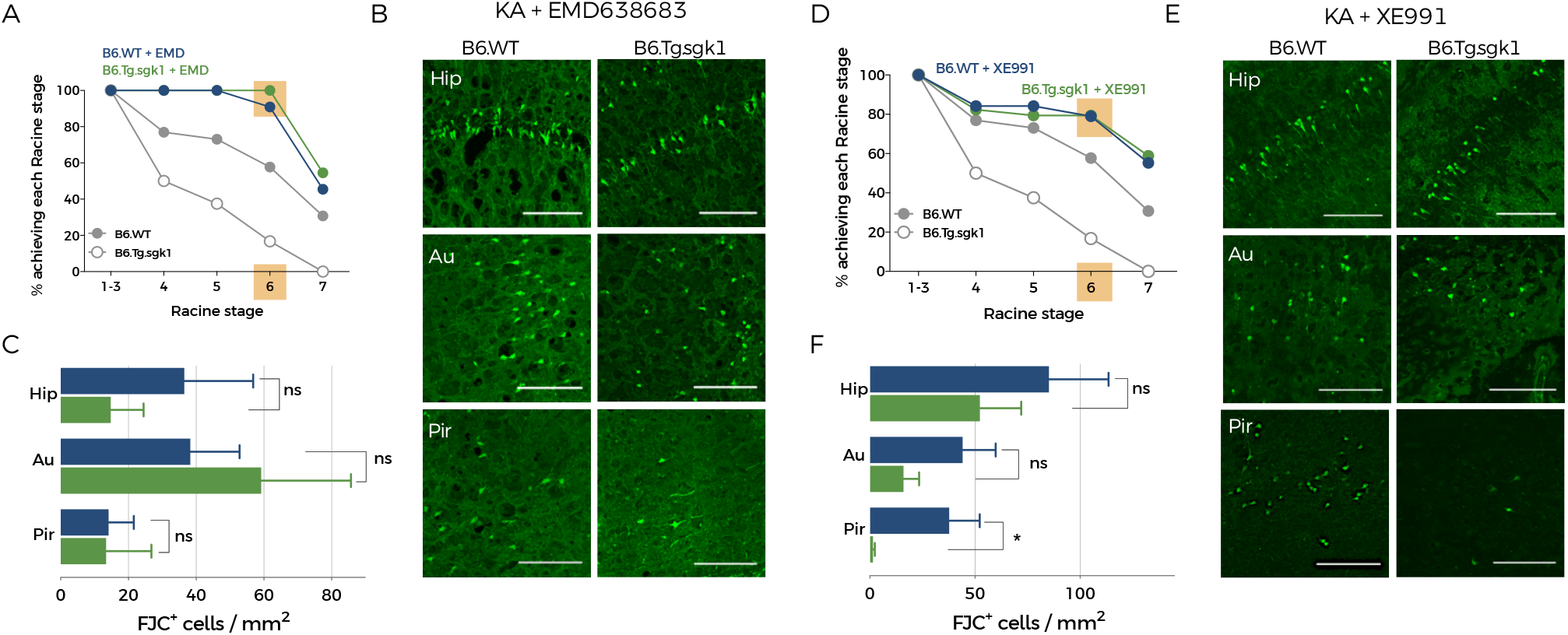
SGK1.1 has a dual role as an anticonvulsant and as a neuroprotective factor in SE. (A) Cumulative plot representing the percentage of EMD638683-pretreated B6.WT and B6.Tg.sgk1 mice reaching the indicated Racine stages after KA injection. Mice reaching stage 6 were used to evaluate neuronal death (highlighted in orange). Data from EMD638683-untreated animals (Figure 1A) are shown in grey for reference. (B6.WT N=26, B6.Tg.sgk1 N=21). (B) Representative confocal images corresponding to FJC staining of brain sections from B6.WT (left column) and B6.Tg.sgk1 mice (right column) pretreated with EMD638683. Brain regions shown are hippocampus (Hip), auditive cortex (Au) and piriform cortex (Pir). Scale bar = 100 μm. (C) Quantitative analysis of FJC-positive cells in EMD638683-pretreated B6.WT (N=8; n=40) and B6.Tg.sgk1 (N=5; n=25) mice in the areas shown in B. Data are mean ± SEM (Mann-Whitney test; ns, not significant). (D) Cumulative plot representing the percentage of XE991-pretreated B6.WT and B6.Tg.sgk1 mice reaching the indicated Racine stages after KA injection (B6.WT N=26, B6.Tg.sgk1 N=24). Mice reaching stage 6 were used to evaluate neuronal death (highlighted in orange). Data from XE991-untreated animals (Figure 1A) are shown in grey for reference. Part of the data in this graph have been obtained from (Armas-Capote et al., 2019). (E) Representative images of FJC stained brain sections from animals in A corresponding to hippocampus (Hip), auditive cortex (Au) and piriform cortex (Pir). Scale bar = 100 μm. (F) Quantitative analysis of FJC-positive cells on brain slices from B6.WT and B6.Tg.sgk1 mice in the areas shown in E. Data are mean ± SEM (N=7, n=35; Mann-Whitney test; *p>0.05; ns, not significant).

We have previously demonstrated that the regulation of the M-current by SGK1.1 underlies the protection mechanism against seizures progression and severity, since pre-treatment of mice with XE991 fully counteracts the effect of the kinase in transgenic mice (Armas-Capote et al. 2019 and Fig.2D). Quantification of neuronal death in this experimental condition allowed us to assess the existence of neuroprotective mechanisms that are independent of the alterations in neuronal excitability associated to SGK1.1 activation. Augmented cell excitability in XE-991 transgenic neurons led to less pronounced differences in neuronal loss between WT and Tg.sgk1 genotypes (Figure 2D-F). However, we still observed a statistically significant reduction in neuronal death levels at the piriform cortex from Tg.sgk1 mice, and a clear tendency towards decreased death levels in all the other brain areas studied (Figure 2F). Importantly, piriform, perirhinal and entorhinal cortex have been described as key seizure-trigger zones (Vismer et al., 2015). This result strongly supports the hypothesis that SGK1.1 neuroprotective role may include additional mechanisms that are independent of its effects on neuronal excitability.

### Activation of sgk1.1 reduces levels of reactive gliosis after status epilepticus

Glial cells have an essential role in maintaining brain homeostasis. Growing evidence supports the hypothesis that inflammation is a consequence as well as a cause of epilepsy (Vezzani, 2007; Choi et al., 2009; Riazi et al., 2010; Vezzani et al., 2011). We quantified the occurrence of gliosis on brain slices from WT and Tg.sgk1 after SE (Racine stage 6). First, GFAP was used as a marker of astrocyte activation to assess astrogliosis levels. As shown in Figure 3A-B, SGK1.1 activation in Tg.sgk1 mice significantly reduced astrogliosis secondary to KA injection in all brain areas, including hippocampus and cortex. Similar to previous observations, this effect was retained in FVB mice, indicating that it was not associated to specific genetic backgrounds (Figure 3C-D).

**Figure 3.**
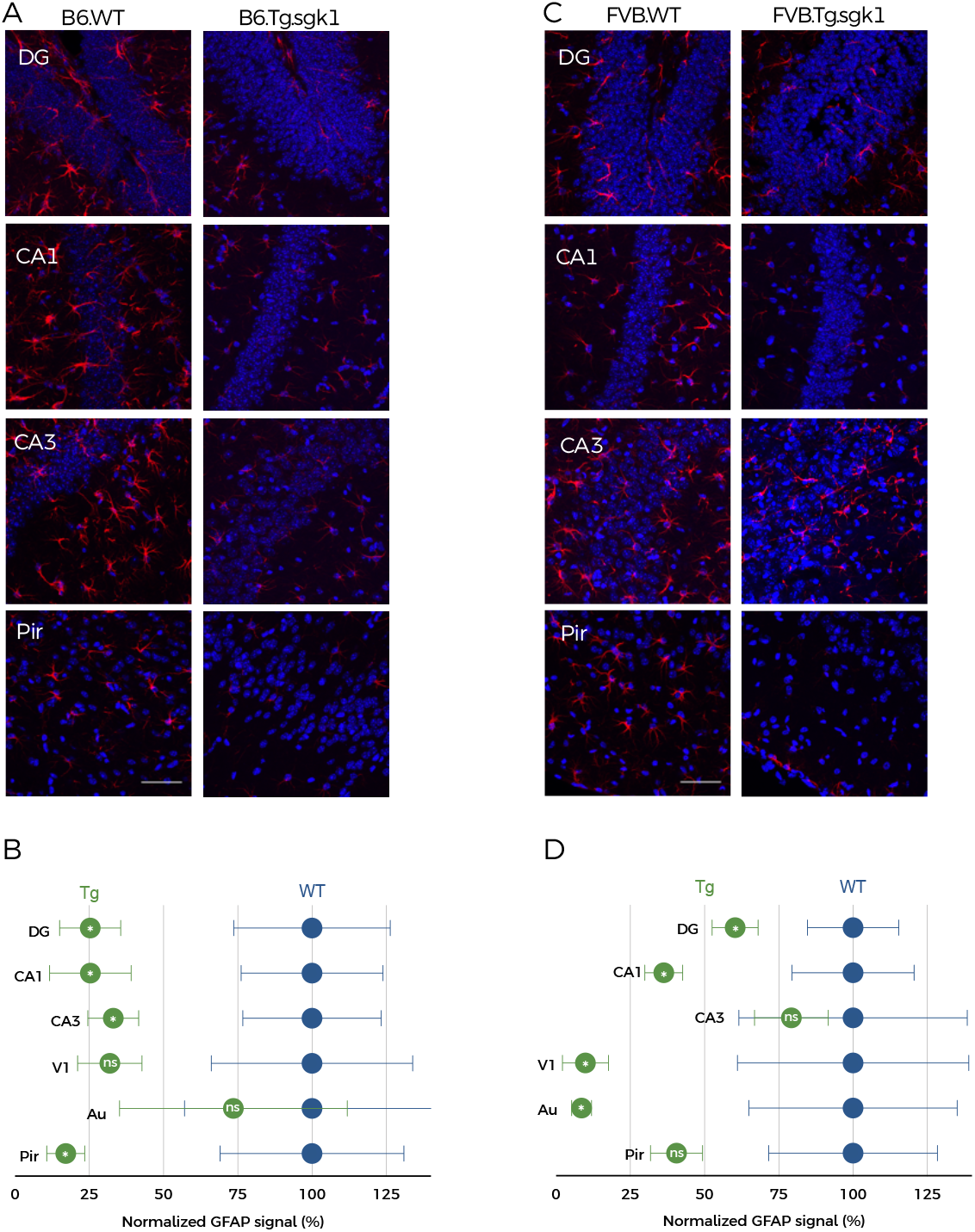
Astrogliosis is decreased in hippocampus and cortex of KA-injected Tg.sgk1 mice. (A) Representative confocal images showing GFAP immunostaining (red, Alexa 494) and DAPI-stained nuclei (blue) for B6.WT and B6.Tg.sgk1, 72h after KA injection in hippocampus (regions DG, CA1 and CA3) and piriform cortex (Pir). Scale bar = 50 μm. (B) Average GFAP levels in transgenic mice compared to WT. Data are mean ± SEM, normalized to WT levels (N=5; multiple t-test; *p<0,05; ns, not significant). (C) Representative images showing GFAP-positive astrocytes in FVB.WT and FVB.Tg.sgk1 mice, 72h after KA injection in same regions as A. (D) Average GFAP levels in transgenic mice compared to WT. Data are mean ± SEM normalized to WT levels (N=6; multiple t-test; *p<0,05; **p<0,01; ns, not significant).

Additionally, we measured microgliosis levels using Iba-1 as a marker. Quantitative analysis revealed significantly lower levels of reactive microglia in B6.Tg.sgk1 mice compared to B6.WT (Figure 4A-B). Similar results were obtained after evaluating the levels of reactive microglia in FVB.WT and FVB.Tg.sgk1 mice, indicating that the effects of SGK1.1 activation on microgliosis are also independent of the genetic background (Figure 4C-D). Our results unveil a new role of SGK1.1 in diminishing reactivity of glial cells, potentially leading to reduced levels of inflammation and structural changes in the damaged brain.

**Figure 4.**
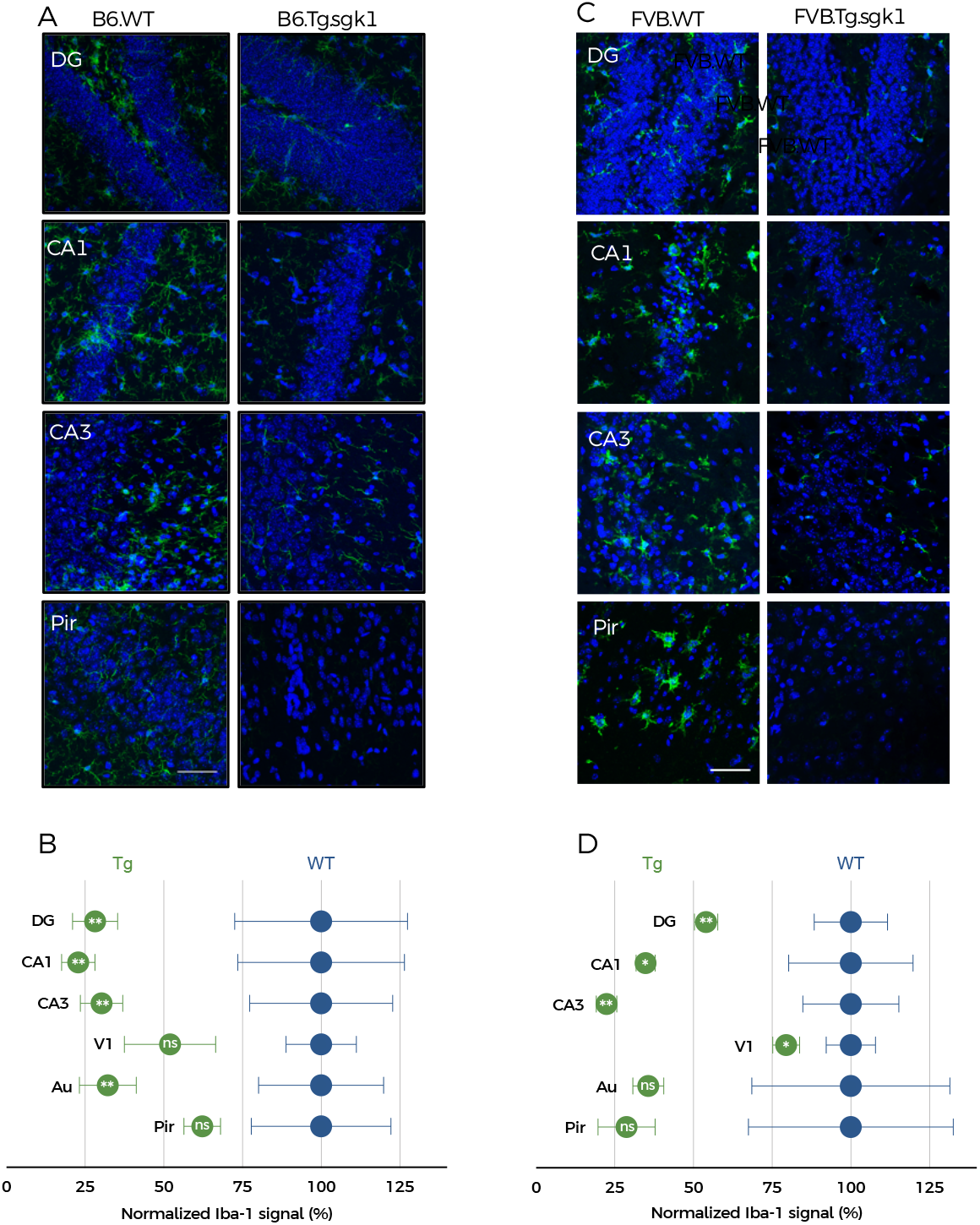
Microgliosis levels are reduced in Tg.sgk1 *vs.* WT mice after KA treatment. (A) Representative confocal images showing Iba-1 reactivity (green, Alexa 488) and DAPI-stained nuclei (blue) in brain slices from B6.WT and B6.Tg.sgk1 mice after KA injection in hippocampus regions (DG, CA1 and CA3) and piriform cortex (Pir). Scale bar = 50 μm. (B) Average Iba-1 levels in transgenic mice compared to WT. Data are mean ± SEM normalized to WT (N=5; multiple t-test; *p<0,05; ns, not significant). (C) Representative confocal images showing Iba-1 staining in FVB.WT and FVB.Tg.sgk1 brain slices after KA injection in same areas as A. (D) Average Iba-1 levels in transgenic mice compared to WT. Data are mean ± SEM normalized to WT (N=5; multiple t-test; *p<0,05; ns, not significant).

### SGK1.1 is mainly expressed in pyramidal neurons

The distribution of the SGK1.1 protein in the brain remained elusive, due to the lack of isoform-specific antibodies. To overcome this limitation, we developed a novel rabbit polyclonal antibody raised against an isoform-specific epitope purified from *E. coli*. The new serum specifically recognized SGK1.1 without cross-reacting with the endogenously expressed ubiquitous isoform SGK1 (Figure 5A). Immunohistochemical localization of SGK1.1 in mouse hippocampus and cortex revealed specific expression in pyramidal neurons as demonstrated by the co-localization with CaMKII (Figure 5B-C). These results are consistent with our previous report that SGK1.1 mRNA is preferentially expressed in pyramidal neurons of the cortex and hippocampus (Wesch et al., 2010). Interestingly, in hippocampus we observed homogeneous expression in all regions that constitute the glutamatergic circuit involved in excitatory synaptic transmission (see upper panels in Figure 5B - CA1- and 5E -CA3-). Double immunohistochemistry also revealed that SGK1.1 is not expressed in parvalbumin-positive interneurons (Figure 5D) or GFAP-expressing astrocytes (Figure 5E-F), further reinforcing the idea that expression of this kinase is restricted to pyramidal neurons, as previously suggested by *in situ* hybridization (Wesch et al., 2010). Within pyramidal neurons, expression is detectable in the soma and neuronal processes (Figure 5B and Figure 5E, left panels), with no positive signal in the nuclei. This localization pattern strongly suggests that SGK1.1 site of action is primarily neuronal and that the decrease in reactive gliosis is secondary to the reduction of neuronal death.

**Figure 5.**
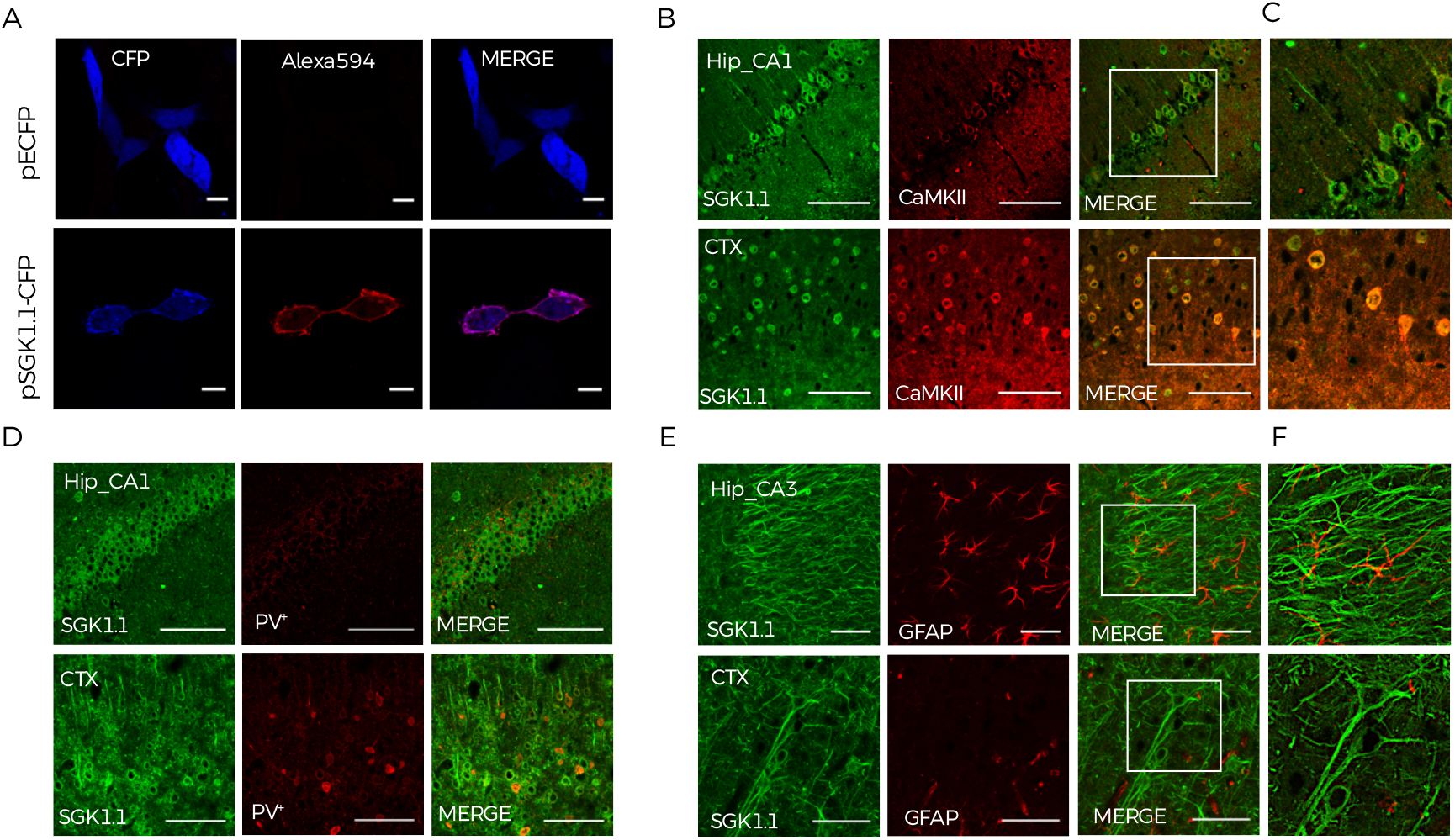
SGK1.1 is selectively expressed at the soma and processes of pyramidal neurons. (A) Representative confocal images of HEK293T cells transfected with an empty vector (peCFP-N1) or a vector expressing SGK1.1-CFP and stained with anti-SGK1.1 rabbit polyclonal antibody. Scale bar = 10 μm. (B) Representative confocal images of WT brain slices showing SGK1.1 (green, Alexa 488) and CaMKII (red, Alexa 594) immunostaining of hippocampus CA1 (Hip_CA1) and visual cortex (CTX) areas. Scale bar = 100 μm. (C) Magnification of white-framed regions in B. (D) Representative confocal images showing SGK1.1 (green) and PV (red) immunostaining of hippocampus CA1 (Hip_CA1) and visual cortex (CTX) from WT brain slices. (E) Representative confocal images showing SGK1.1 (green, Alexa 488) and GFAP (red, Alexa 594) expression of hippocampus CA3 (Hip_CA3) and visual cortex (CTX). (F) Magnification of white-boxed areas in E.

### SGK1.1 decreases apoptosis after KA-induced SE

Given the fact that SGK1.1 catalytic domain shares high homology with the catalytic domain of the well described anti-apoptotic kinase AKT (Kobayashi et al., 1999), and that the ubiquitous isoform SGK1 has anti-apoptotic activity (Ferrelli et al., 2015), we evaluated the ability of SGK1.1 to reduce the levels of apoptosis in an *in vitro* model using the TUNEL assay (Crowley et al., 2016). As previously described, apoptosis levels were significantly reduced in H_2_O_2_-treated HEK293T cells expressing AKT (Park et al., 2016) (Figure 6A, middle row and Figure 6B, grey bar). Heterologous expression of the constitutively active form of SGK1.1 lead to similar results, supporting the potential role of this kinase as an antiapoptotic factor (Figure 6A-lower row and Figure 6B, green bar). In contrast, this effect was abolished in cells expressing a kinase-dead mutant of SGK1.1 (K220A) (Figure 6B, orange bar) (Arteaga et al., 2008; Wesch et al., 2010). Finally, heterologous expression of a mutant with constitutive nuclear localization (FF19.20AA) showed higher sensitivity to H_2_O_2_-induced apoptosis (Figure 6B, purple bar), indicating that correct subcellular targeting is essential for SGK1.1 anti-apoptotic effects.

**Figure 6.**
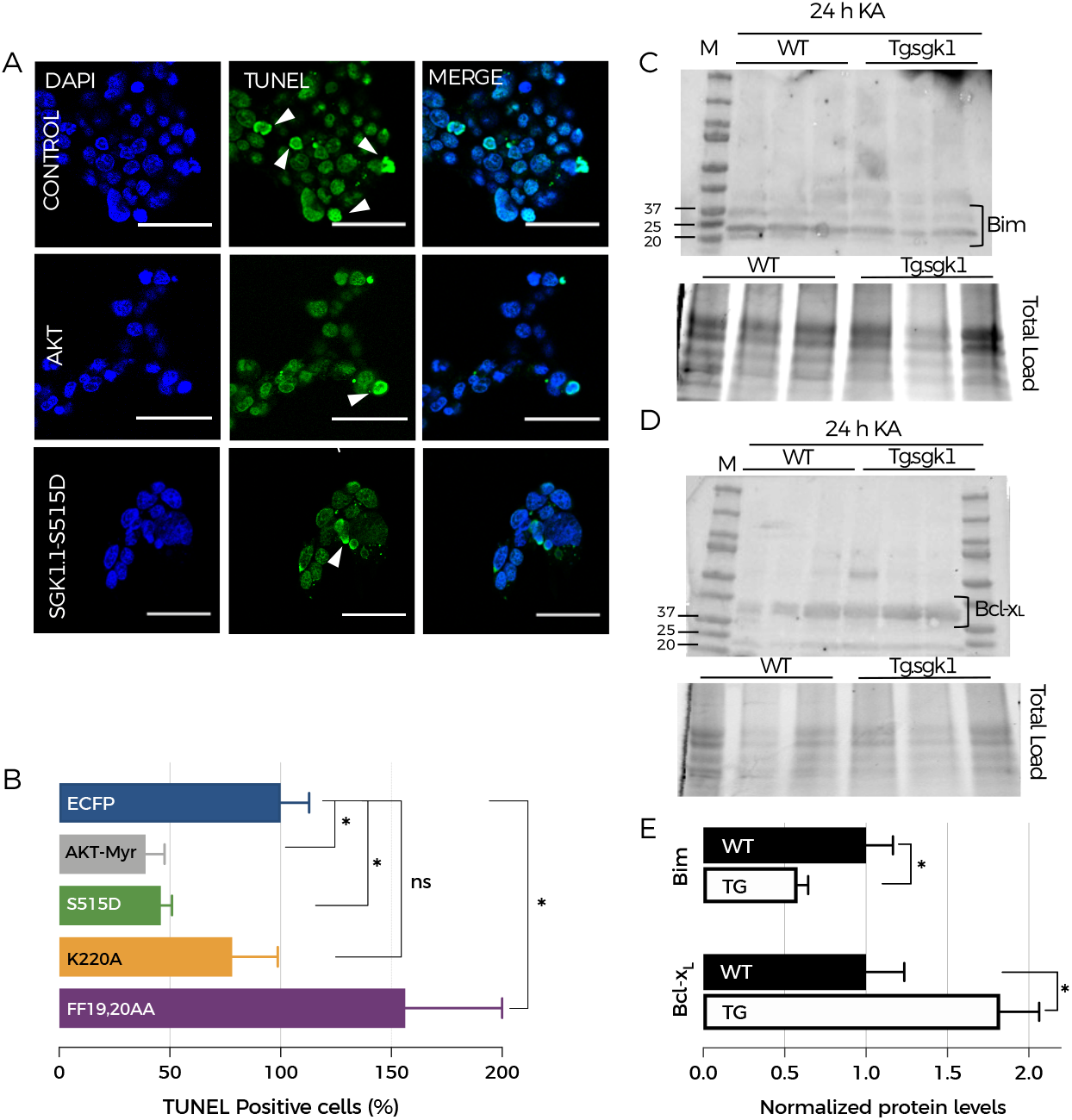
SGK1.1 activation exerts anti-apoptotic effects via modulation of apoptosis mediators after KA-induced SE. (A) Representative confocal images of fluorescein-stained cells transfected with the indicated constructs. Arrows point to TUNEL-positive cells. Scale bar = 10 μm. (B) Quantitative analysis of TUNEL-positive cells after transfection with the indicated constructs and H_2_O_2_ treatment. Bars show mean ± SEM (n=5; cells were counted on at least 6 different fields per condition; one-way ANOVA followed by multiple comparison test; * p<0.05; ns, not significant). (C) Top, Representative western blot of Bim protein abundance in the hippocampus of WT and transgenic mice 24h after KA administration. Bottom, total protein loading. (D) Top, Representative western blot of Bcl-x_L_ protein abundance in the hippocampus of WT and transgenic mice 24h after KA administration. Bottom, total protein loading. (E) Quantification of western blot data. Bars represent mean ± SEM from WT (n=4) and Tg.sgk1 (n=6; Unpaired t-test; *p<0.05). Total protein fluorescence staining (stain-free™, BioRad) was used as loading control.

Altogether, these results support the idea that SGK1.1 could act as an anti-apoptotic factor *in vivo*, decreasing SE-induced apoptosis in the mouse hippocampus. We tested this hypothesis by quantifying the expression of different markers of apoptotic pathways. First, we evaluated how activation of SGK1.1 in the transgenic mice affected the levels of Bim, a known pro-apoptotic marker of the intrinsic apoptosis pathway (O’Connor et al., 1998; Engel and Henshall, 2009; Kim et al., 2014). Western blot analysis performed 24 h after KA-induced SE showed significantly decreased levels of Bim in Tg.sgk1 mice *vs.* WT (Figure 6C and Figure 6E). Bands corresponding to the molecular mass of the three existing splice isoforms of Bim (Bim_s_, Bim_L_ and Bim_EL_) (Putcha et al., 2001) were detected (Figure 6C). Additionally, we analyzed the expression of Bcl-x_L_, a member of the anti-apoptotic Bcl-2 protein family. This factor has been proposed to reduce apoptosis by preventing mitochondrial permeability transition associated to release of pro-apoptotic factors, therefore contributing to maintain cell viability in the central nervous system (Boise et al., 1993; Krajewska et al., 2002; Jonas et al., 2014). Consistent with the hypothesis that SGK1.1 activation can exert anti-apoptotic effects, western blot experiments show significantly increased Bcl-x_L_ protein abundance in the hippocampus of Tg.sgk1 mice *vs.* WT following KA-induced seizures (Figure 6D and Figure 6E).

Taken together, our results show that activation of SGK1.1 is a potent mechanism of neuroprotection after SE, with a striking reduction in neuronal death accompanied by a very prominent reduction in reactive gliosis. Mechanistically, this effect underlies a combination of two main factors. In addition to previously demonstrated decreased neuronal excitability due to increased M-current density, we now demonstrate that SGK1.1 activation triggers anti-apoptotic signaling pathways involving altered levels of Bim and Bcl-x_L_ proteins.

### Transgenic mice do not show hippocampal ectopic neurogenesis

The existence of aberrant hippocampal neurogenesis in the subgranular region of the DG has been related to epilepsy (Parent et al., 2006; Chugh et al., 2015). Previous studies have suggested that SGK1 might be associated with neurogenesis in hippocampus triggered by glucocorticoid hormones (Anacker et al., 2013). Therefore, if SGK1.1 is to be evaluated as a valid pharmacological target in the treatment of epilepsy, it is important to determine whether constitutive activation of SGK1.1 in our animal model has side effects on ectopic neurogenesis. In this study, we quantified neurogenesis by BrdU incorporation and DCX staining in granular cells of the DG and hilus (Figure 7). Our results showed no significant differences in normal or ectopic neurogenesis in transgenic mice *vs.* WT (Figure 7). This finding supports the conclusion that activation of SGK1.1 does not produce aberrant neurogenesis.

**Figure 7.**
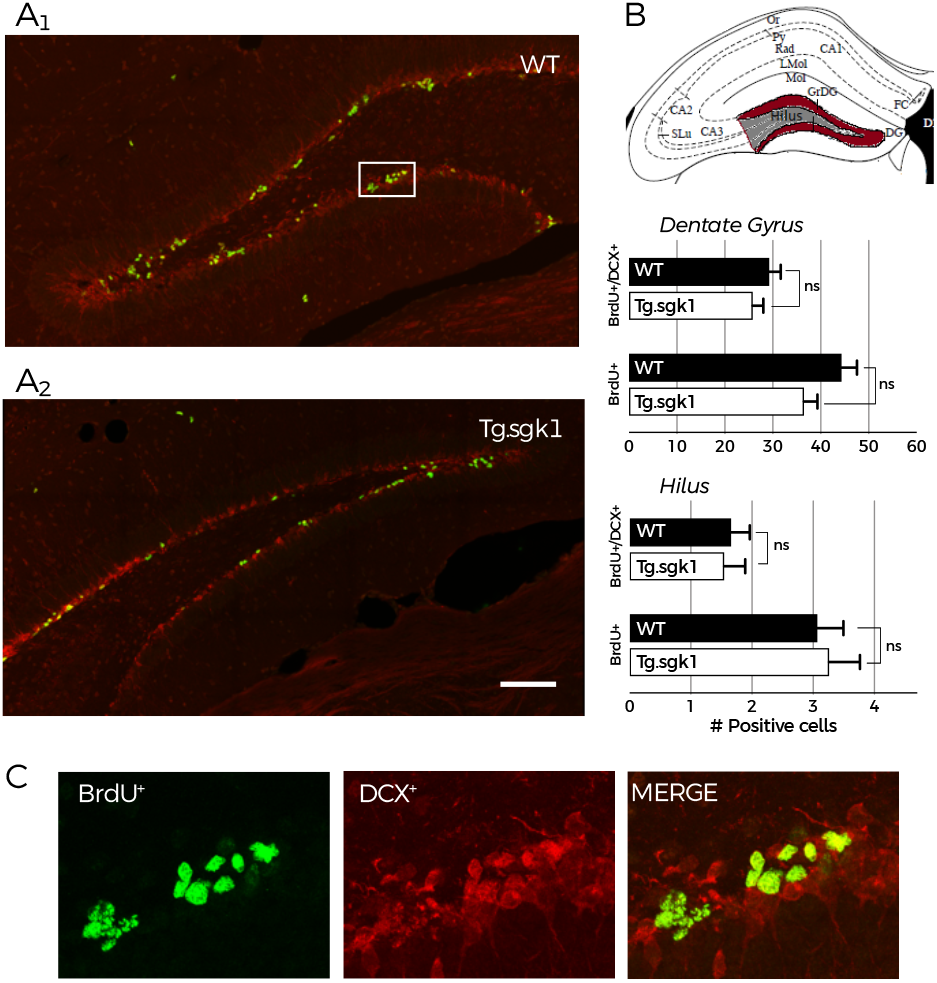
Activation of SGK1.1 does not alter neurogenesis in hippocampus. (A) Representative confocal images showing merged immunostaining signals corresponding to BrdU (green) and DCX (red) from B6.WT (A_1_) and B6.Tg.sgk1 (A_2_) in hippocampus DG. Scale bar= 100 μm. (B) *Top*, schematic representation showing brain areas for neurogenesis quantification. *Bottom*, quantification of BrdU^+^ and DCX^+^ cells per brain section in granular cells of DG (newborn cells, *above*) and hilus (ectopic neurogenesis, *below*). Bars represent mean ± SEM (N=4; unpaired t-test; ns, not significant) (C) Magnification of white-boxed area in A1. BrdU^+^ cells (green), DXC^+^ cells (red) and merged image.

## DISCUSSION

Our work demonstrates that SGK1.1, which has been previously shown as a modulator of Kv7 channels that potently reduces KA-induced seizure severity (Miranda et al., 2013; Armas-Capote et al., 2019), exerts a dual protecting role by additionally modulating and reducing seizure-induced cell damage in the brain. Importantly, this effect is robust and independent of the genetic background. It is maintained in FVB/NJ mice, a strain associated to higher levels of neurodegeneration than C57BL/6J (Royle et al., 1999; Kasugai et al., 2007).

Neuroprotection is due to SGK1.1 activation, since acute kinase inhibition with the specific inhibitor EMD638683 lead to equal levels of neuronal death in both genotypes. This result also rules out that the observed effects are a consequence of long-term changes induced by the expression of the transgene. Inhibition of the M-current by pre-treatment with the Kv7 blocker XE991 abolished differences in behavioral seizures between genotypes (Fig.2), as well as the decrease in CA1 excitability associated to activation of the kinase (Armas-Capote et al., 2019). Strikingly, transgenic mice still showed significantly reduced neuronal death levels in relevant brain areas after the KA challenge, suggesting that activation of SGK1.1 modulates pathways preventing neuronal death independently of its role in membrane Kv7 modulation.

The mechanism underlying this observed SGK1.1-mediated protection against neuronal death relays at least partially on the role of this kinase as an anti-apoptotic factor in the brain. First, our data show that SGK1.1 activation reduces H_2_O_2_-induced apoptosis to similar extents than AKT. This finding is not surprising, given the shared homology of SGK1.1 and AKT catalytic domains. Further, the existence of such homology makes it tempting to hypothesize that both kinases might share downstream substrates including the anti-apoptotic transcriptional factor Bim, which has been related to SE-brain damage with FOXO3a and caspase-3 (Kim et al., 2014). Consistent with this notion, we now show that activation of SGK1.1 in Tg.sgk1 mice reduces Bim protein levels in hippocampus, whereas Bcl-x_L_ is upregulated. Importantly, both members of the Bcl-2 family have been described as key regulators of the mitochondrial (intrinsic) apoptotic pathway (Youle and Strasser, 2008) and related to seizure-induced brain damage (Murphy et al., 2010; Kim et al., 2014). Increased levels of Bim have been reported in mice and rats after KA-induced SE, while Bim-deficient mice show reduced levels of neurodegeneration (Motoyama et al., 1995). Bim localizes to intracellular membranes, where it has been proposed to induce apoptotic cell death via caspase activation (O’Connor et al., 1998). This proapoptotic effect is blocked by interaction with Bcl-xL, an anti-apoptotic factor found in mature neurons in the adult brain (Motoyama et al., 1995; Roth et al., 2000). Accordingly, Bcl-x_L_-deficient mice show massive neuronal cell death (Motoyama et al., 1995). Our results showing modulation of both factors in transgenic mice are consistent with the observed significantly lower levels of neurodegeneration that are associated to activation of SGK1.1.

A potential ramification of the anti-apoptotic role of SGK1.1 could be the existence of deleterious pro-proliferative effects (Brunet et al., 2001). In the framework of the observed neuroprotective effects of SGK1.1 activation, and taken into account previous reports of SGK1 effects (Anacker et al., 2013), we were especially concerned about the potential induction of aberrant hippocampal neurogenesis in the DG (Parent et al., 2006; Löscher, 2012; Chugh et al., 2015). It has been proposed that this ectopic neurogenesis after seizure-induced damage could result in the alteration of neuronal electrophysiological properties, contributing to the generation of hyperexcitable circuits. This process has been involved in the maintenance of epileptic activity (Ribak et al., 2000; Parent et al., 2006). However, our data did not reveal significant differences in the levels of ectopic neurogenesis in the DG of Tg.sgk1 *vs.* WT mice, ruling out this possibility (Parent et al., 2006; Anacker et al., 2013). It will be interesting to address in the future whether there is any difference in ectopic neurogenesis between Tg.sgk1 and WT mice after KA treatment, both in acute conditions as well as in animals undergoing recurrent spontaneous seizures (acquired epilepsy). Neurodegeneration has been described as one of the most common alterations found in epilepsy patients contributing to progression towards a sclerotic and hardened brain (Mathern et al., 1997; Mathern et al., 2002; Goldberg and Coulter, 2013). Although there is a growing body of evidence supporting that neuroprotection does not prevent development of acquired epilepsy (André et al., 2001; Löscher, 2002; Pitkänen, 2002; Brandt et al., 2003), it has been proven that alleviation of neurodegeneration diminishes adverse effects of SE and might improve recovery (Bolanos et al., 1998; Pitkänen and Kubova, 2004; Brandt et al., 2006). Proinflammatory mediators, reactive astrocytes and microglia have been found in the resected hippocampi of TLE patients and might contribute to the generation of new seizures (Crespel et al., 2002; Aronica et al., 2007; Ravizza et al., 2008; Van Gassen et al., 2008). However, it is not clear whether inflammation is a consequence or a cause of epilepsy (Vezzani, 2007; Choi et al., 2009; Riazi et al., 2010; Vezzani et al., 2011). In the present work, we demonstrate that while SGK1.1 is not expressed in astrocytes, its neuronal activation prevents reactive gliosis in a transgenic mouse model, limiting the extent of overall brain damage. Furthermore, our results support the idea that, at least in the animal model used in this study, gliosis occurs as a consequence of neuronal death.

Although epilepsy research has allowed the development and improvement of different available treatments, there is still a great need for approaches able to control seizures in approximately 30% of the patients showing resistance to antiepileptic drugs (Löscher and Schmidt, 2011). New strategies suitable to diminish and reduce brain damage after SE might help to prevent the progression of the disease as well as the deterioration of neuronal tissue, which is usually accompanied by a long list of comorbidities and psychiatric dysfunctions (Kobau et al., 2006; Tellez-Zenteno et al., 2007; Elliott et al., 2009; Kanner et al., 2010). The present work, together with our previously published results (Miranda et al., 2013; Armas-Capote et al., 2019) strongly supports a dual role of SGK1.1 as an anti-convulsant and neuroprotective factor, which could open new therapeutic avenues in epilepsy.

## Acknowledgments

We thank Dr Tomás Gonzalez and Dr Pedro Barroso for the help with FJC protocol. T.G. and D.A.R are members of the Red de Excelencia “Iniciativa Española en Canales Iónicos”.

